# Dominance vs. epistasis: the biophysical origins and plasticity of genetic interactions within and between alleles

**DOI:** 10.1101/2022.04.03.486855

**Authors:** Xuan Xie, Ben Lehner, Xianghua Li

## Abstract

A central challenge in genetics, evolutionary biology and biotechnology is to understand and predict how mutations combine to alter phenotypes, including molecular activities, fitness and disease. In diploid organisms, two mutations in the same gene can either combine on the same chromosome or on different chromosomes, with interactions between the mutations quantified as epistasis and dominance, respectively. However, a direct comparison of the extent, sign and stability of interactions within and between alleles is lacking. Here we show that, even in the simplest biophysical systems, interactions between mutations are frequent, context-dependent and different when variants are combined within and between alleles. Whereas protein folding alone generates epistasis, the addition of a single molecular interaction is sufficient to cause dominance. Epistasis and dominance interactions change quantitatively, qualitatively and differently as a system becomes more complicated or the conditions change. Altering the concentration of a ligand can, for example, switch an allele from dominant to recessive. Our results show that epistasis and dominance should be widely expected in even the simplest biological systems but also reinforce the view that they are plastic system properties and so a formidable challenge to predict. Accurate prediction of epistasis and dominance will require either detailed mechanistic understanding and experimental parameterization or brute-force measurement and learning.

## Introduction

A fundamental goal in biology is to understand and predict how mutations combine to alter phenotypes. This is important in biotechnology – for example when engineering new enzymatic activities and protein properties – and also in animal and plant breeding, clinical genetics and evolutionary biology.

Although mutations are normally assumed to have independent effects, this often proves not to be the case: additional variants within the same gene as well as in other genes can quantitatively and qualitatively alter the impact of a mutation (Domingo et al., 2019). Predicting these genetic interactions between variants and so improving genetic prediction beyond the performance that can be achieved using additive models is a central challenge in clinical genetics, evolutionary biology, agriculture and biotechnology.

When two variants occur in the same gene in a diploid species, they can either both occur on the same chromosome or each variant can be on a different chromosome, i.e., as on the paternal and maternal alleles. In any individual, approximately 1 in 10 human genes carry two or more variants compared to a reference genome (Frazer et al., 2020), with >20,000 combinations of variants within the same gene observed >1.5 million times in 59 medically actionable genes in ∼50k individuals from the UK Biobank population (Van Hout et al., 2020). It is therefore important for clinical genetics to understand and be able to predict what happens when multiple mutations occur in the same gene, and how the outcome differs depending upon whether the two variants occur on the same chromosome (‘within allele’) or on different chromosomes (‘between alleles’).

In quantitative genetics, clinical genetics, functional genomics, animal breeding and evolutionary biology, the phenotypic change when combining mutations is most often assumed to be additive or log-additive (Domingo et al., 2019; Mani et al., 2008). For example, when two variants are combined within the same copy of a gene, the change in phenotype or fitness is often expected to be log-additive, with deviance from this expectation referred to as a genetic interaction or epistasis (Domingo et al., 2019; Phillips, 2008). Similarly, when two variants are combined in different alleles (i.e. as a compound heterozygote), the expected phenotype is normally considered to be the average of the phenotypes when the variants are present in two copies (i.e. homozygotes), with any deviance from this additive expectation quantified as dominance (Falconer and Falconer, 1989; Kacser and Burns, 1981; Wright, 1934). Mutations have been found to vary extensively in their dominance, with important implications for breeding, evolution and human clinical genetics (Amberger et al., 2019; Veitia et al., 2018).

Various mechanisms have been proposed to cause epistasis, dominance, or both (Domingo et al., 2019; Keightley, 1996; Omholt et al., 2000; Veitia et al., 2018). These include the non-linear relationships between additive biophysical parameters and phenotypes, for example, the sigmoidal relationship between the folding energy of a protein and the fraction of a protein that is folded (Olson et al., 2014; Otwinowski et al., 2018; Tokuriki et al., 2007; Wylie and Shakhnovich, 2011). Additional non-linear relationships between genotype and phenotype are introduced by molecular interactions (Diss and Lehner, 2018; Otwinowski, 2018), cooperativity (Boj et al., 2010; McKeown et al., 2014), molecular competition (Baeza-Centurion et al., 2019; Porter et al., 2017), metabolic flux (Kacser and Burns, 1981; Lunzer et al., 2005) and feedback loops and other dynamics in cellular networks (Omholt et al., 2000). In addition to such ‘global’ or ‘non-specific’ interactions between mutations due to nonlinear genotype-phenotype relationships (Domingo et al., 2019), mutation-specific causes of epistasis and dominance are also observed. These include non-additive changes in free energy when mutating energetically-coupled residues (Horovitz et al., 2019) and gain-of-function dominant negative mutations (Wilkie, 1994).

Protein (or RNA) folding and binding to ligands constitute the fundamental reactions common to nearly all cellular processes. However, how these foundational biophysical processes cause interactions between mutations, and how these interactions differ when mutations are combined within the same allele (epistasis) versus between two alleles (dominance) has not been comprehensively explored. Here we address this shortcoming by quantifying the interactions between mutations in statistical thermodynamic models of protein folding and binding. Thermodynamic models have previously proven useful for interpreting and predicting how mutations combine in large-scale experimental datasets (Adams et al., 2016; Domingo et al., 2019; Faure et al., 2022; Kinney et al., 2010; Li et al., 2019; Li and Lehner, 2020; Otwinowski, 2018).

We show that, even in the simplest biophysical systems, mutations should be expected to frequently combine with outcomes that differ from the additive or log-additive expectations. Moreover, the expected outcome is typically different when combining variants within versus between alleles. In the simplest possible protein system, where a phenotype is linearly dependent on the concentration of a folded protein, there is no dominance (between-allele interactions) but abundant epistasis (within-allele interactions). Adding a single ligand-binding reaction to the system is sufficient to generate dominance, with dominance changing both quantitatively and qualitatively depending on how much ligand is present. Furthermore, in this simple biophysical system, the epistasis depends on which biophysical parameter (folding or binding energy) is affected by a mutation, whereas the dominance does not differ for mutations altering folding or binding. Introducing a nonlinear dependency of a phenotype on the concentration of a protein or protein-ligand complex reshapes both dominance and epistasis and often in opposite directions. Our results show that dominance and epistasis are expected in even the simplest biophysical systems but that they are plastic with their magnitude and sign dependent on the precise molecular details of the system and also on the conditions. Moreover, they highlight that both types of interaction are difficult to quantitatively and qualitatively predict. Prediction of dominance and epistasis will therefore require either detailed mechanistic understanding and quantification of relevant cellular parameters or large-scale empirical measurements and learning.

## Results

### Within- and between-allele genetic interactions in simple biophysical systems

In a diploid system, each gene is present in two copies – the maternal and paternal allele, respectively. When a gene carries two different mutations the two variants can, therefore, either both be present in the same allele or each variant can be in a different allele (a compound heterozygote, Figure 1a). Even in a haploid organism, a similar comparison can be made for gene duplicates – two variants can be combined in the same duplicate (paralog) or each can be in a different paralog.

**Figure. 1.**
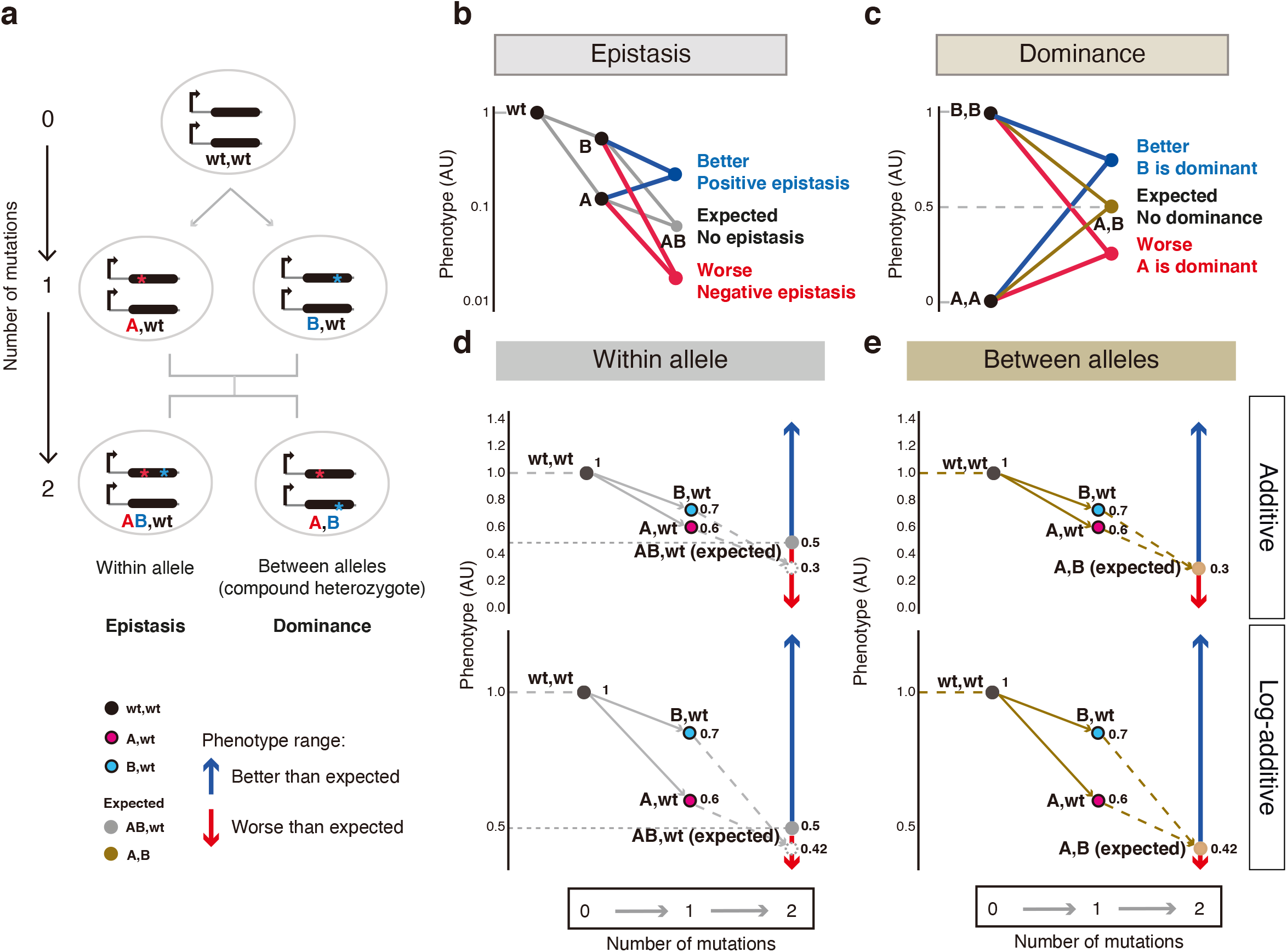
Quantifying within- and between-allele genetic interactions. (a) In diploid systems, two mutations in the same gene can occur in the same allele or one in each allele. (b, c) Epistasis (b) and dominance (c) quantification. (d, e) The scheme of quantifying how two mutations combine within-(d) or between-allele (e) with corresponding additive (d, e upper) and log-additive (d, e lower) assumptions. For within-allele mutation combinations, the lower bound of the expected phenotype is set to 0.5 AU, with the dotted gray empty circle indicating the expected phenotype below 0.5 AU and the solid gray circle indicating the new expected phenotype set to 0.5 AU.

Quantifying how variants interact when they are combined requires the specification of a null model for independent effects. The interactions between variants within the same allele of a gene are typically referred to as epistasis or genetic interactions with a log-additive (or sometimes additive) null model used as the expected outcome (Figure 1b) (Domingo et al., 2019; Mani et al., 2008). In contrast, interactions between variants in different alleles are typically quantified as dominance, which uses an additive expected outcome (Figure 1c, Supplementary Figure 1) (Falconer and Falconer, 1989; Wright, 1934). Therefore, when comparing how mutations interact within and between alleles of a gene, throughout this manuscript we quantify interactions using both of these null models – additive and log-additive – as well as directly compare the phenotypes of double mutants (Figure 1d, e).

### Protein folding generates epistasis but not dominance

We first considered the simplest biophysical system where a phenotype (or fitness) is linearly related to the concentration of a protein that folds cooperatively and exists in two states: unfolded and folded (Model 1) (Figure 2a, b). Such two-state folding is observed for many small proteins (Malhotra and Udgaonkar, 2016). The probability of the protein being folded (or folded fraction, F) depends on the free energy difference between the two states (the folding energy, ΔG_Folding_) (Figure 1c) and can be calculated using the Boltzmann distribution, F = exp(-ΔG_Folding_ / (RT)) / (1 + exp(-ΔG_Folding_ / (RT))), where R is the gas constant and T is the absolute temperature. Mutations affect the folding energy relative to the wild-type (ΔΔG_Folding_) and changes in free energy are assumed to be additive when combining mutations in the same molecule. We first considered a moderately stable protein (ΔG_Folding_ = −2 kcal per mol) and mutations with a range of effect sizes (ΔΔG_Folding_: −2 to 13 kcal per mol) and combined pairs of mutations either within the same allele or between the two alleles (for details, see Methods). The interaction between each pair of mutations was then quantified as the difference between the calculated (observed) double mutant phenotype and that expected from an additive or log-additive model (Figure 1d-k).

**Figure. 2.**
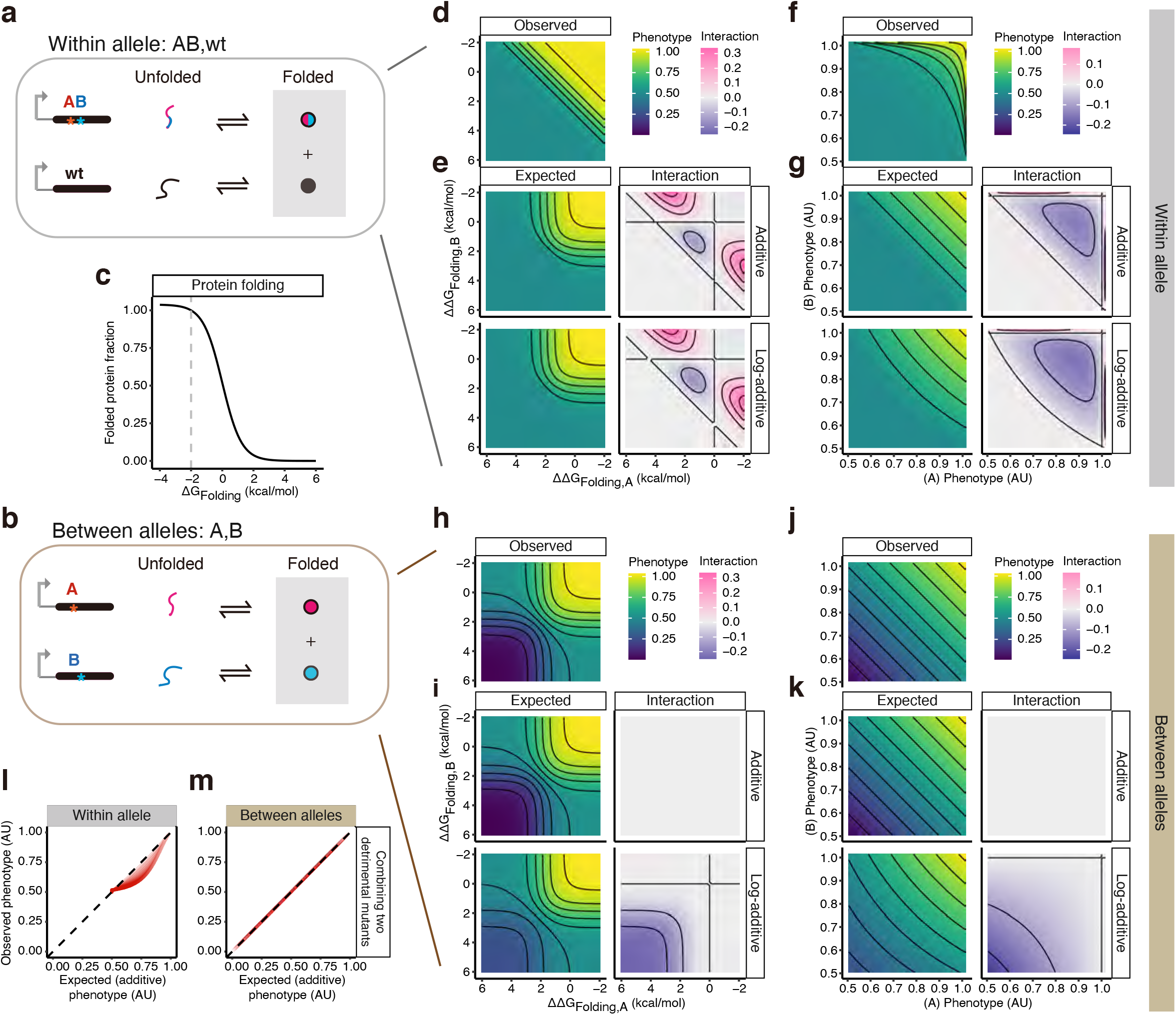
Protein folding generates epistasis but not dominance. (a, b) Two-state protein folding system (Model 1) with two mutations within- (a) or between-alleles (b). Phenotypes are determined by the folded protein concentration marked with grey-shaded boxes. (c) The relationship between the free energy changes of protein folding and folded protein fraction of homozygotes. (d-k) Heatmaps show how two mutations combine within (d-g) or between alleles (h-k) when they are ordered by free energy changes (d, e, h, i) or phenotypes (f, g, j, k). (l, m) Relationships between the observed and expected phenotypes with additive expectation when combining two detrimental mutants within- (l) or between-allele (m).

We plotted the phenotypes of double mutants against the change in free energy (ΔΔG_Folding_) of the constituent single mutants (Figure 2c-e, h, i) and also against the phenotypes of the single mutants (Figure 2f, g, j, k). Comparing the iso-phenotype contour lines (phenotype isochores) when combining mutations within allele (Figure 2d) to the expectations when assuming additivity (Figure 2e upper panel) or log-additivity (Figure 2e lower panel) illustrates how protein folding alone generates epistasis (Domingo et al., 2019; Olson et al., 2014; Tokuriki et al., 2007; Wylie and Shakhnovich, 2011): the sigmoidal relationship between the fraction of a protein that is folded and free energy (Figure 2c) means that two destabilizing mutations often have an outcome that is more detrimental than both the additive and log-additive expectations, and combining a stabilizing with a destabilizing mutation often results in a better than expected outcome (Figure 2e, g, l, Supplementary Figure 2b).

In contrast, when two mutations affecting folding are combined in different alleles, the phenotypic outcome is additive (Figure 2h-k, m), consistent with the most widely used null model for dominance. In comparison, assuming log-additivity overestimates the phenotype when combining two destabilizing mutations, resulting in negative interactions (lower panels of Figure 2i, k). Considering more stable and less stable wild-type proteins does not change these conclusions (Supplementary Figure 2a, c-f).

In summary, protein folding alone is not expected to generate between-allele interactions (dominance) but it does generate within-allele interactions (epistasis): additivity is the correct null model for dominance but neither additivity nor log-additivity are the correct expectations for epistasis.

### Ligand binding generates dominance

Many proteins bind ligands as part of their function, for example small molecules, nucleic acids or other proteins. We examined how ligand-binding affects the expectation for how mutations interact within and between alleles. We first considered the case where a phenotype is linearly determined by the concentration of protein-ligand complex (Model 2) (Figure 3a, b). In this system, mutations can alter the free energy of folding, as in Model 1, or they can alter the binding energy (i.e. affect the binding affinity). In Figure 3c and 3d, we compare the observed and expected double mutant phenotypes when combining mutations affecting either folding or binding in a model where the ligand concentration is the same as the protein concentration, the protein is moderately stable (ΔG_Folding_ = −2 kcal per mol) and the binding affinity is moderate (Jecklin et al., 2009) (free energy of binding ΔG_Binding_ = −5 kcal per mol, corresponding to a dissociation constant, K_D_ = 291 nM at 37 °C).

**Figure. 3.**
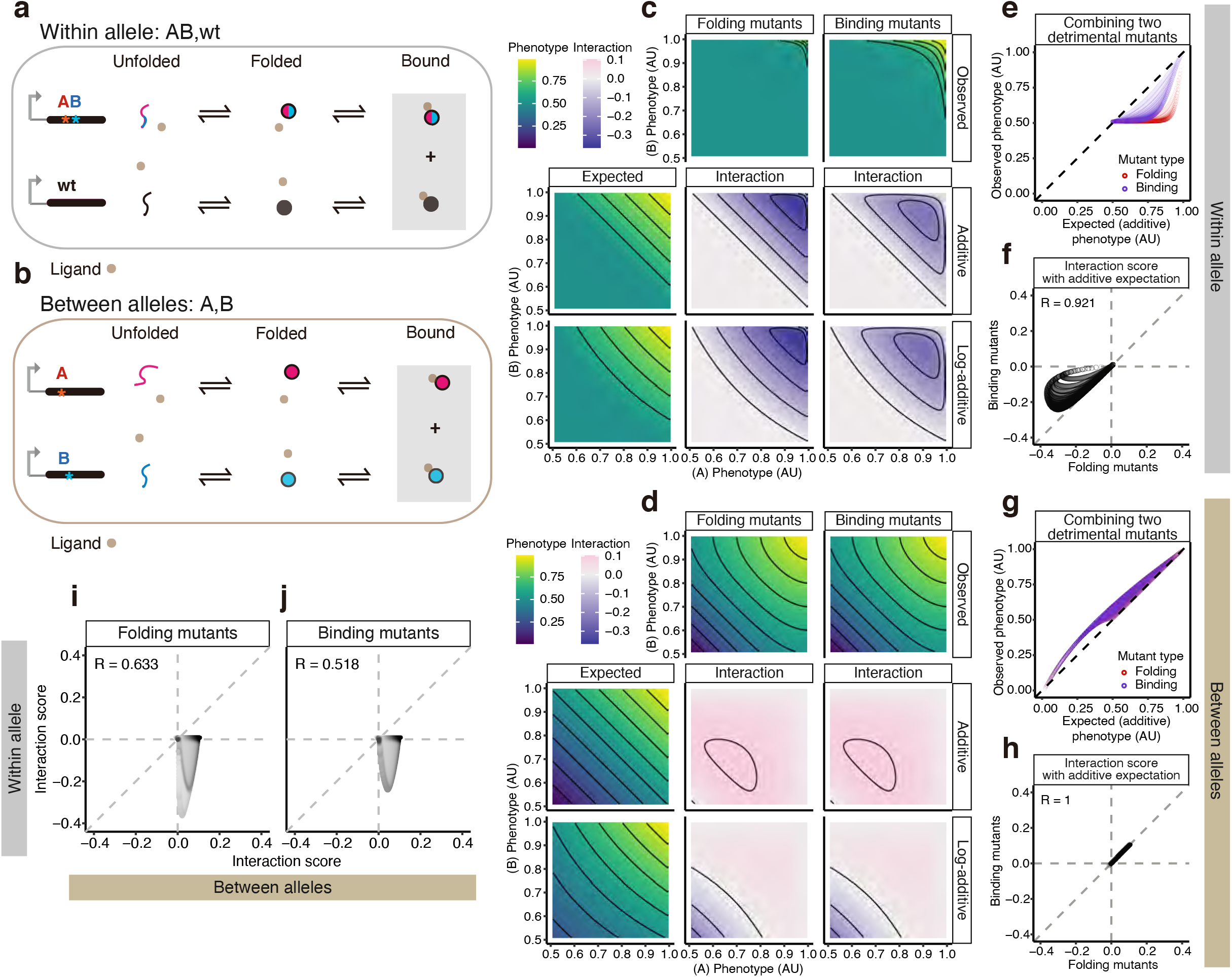
Ligand binding generates dominance. (a, b) Three-state protein system with unfolded, folded, and ligand-bound states (Model 2), with two mutations of the same gene within- (a) or between-alleles (b). Phenotypes are determined by the ligand-bound protein concentration marked with gray-shaded boxes. (c, d) Heatmaps showing how two mutations both affect the same biophysical parameters combine: protein-folding (the second column) or ligand-binding (the third column) when they are ordered by the phenotype. (e, g) Relationships between the observed and expected phenotypes with additive expectation when combining two detrimental mutants within (e) or between allele (g). (f, h, i, j) Comparisons of interaction scores between different types of double mutant combinations: protein-folding vs. ligand-binding mutants within- (f) or between-allele (h), between-allele vs. within-allele interactions of the protein-folding (i) or ligand-binding (j) mutants.

Similar to what is observed in the folding-only model (Model 1), when two destabilizing mutations are combined in the same protein, the change in concentration of the protein bound to the ligand is often larger than the additive or log-additive expectation i.e., there is negative epistasis (Figure 3c and Supplementary Figure 3a, b).

However, unlike in Model 1, there are now non-additive interactions between mutations when they are combined in the two different alleles, with two destabilizing mutations typically having an outcome that is better than the additive expectation (Figure 3d and Supplementary Figure 3c, d). That is, mutations that do not show any dominance in the protein folding-only model now display dominance (Porter et al., 2017).

### Mutations interact with opposite sign within and between alleles

In the folding and binding model (Model 2), mutations interact to generate both epistasis and dominance: the phenotype of double mutants often differs from the additive and log-additive expectations. However, comparing the curvature of the observed phenotypic isochores in Figures 3c and 3d, it can be seen that the interactions within and between alleles are qualitatively different: two variants that reduce stability typically have an outcome that is worse than additive when they both occur in the same allele but better than additive when they occur in different alleles. Moreover, although the signs of interaction are different, the interaction scores when the same mutations are combined within versus between alleles are positively correlated (Figure 3i, j), meaning that mutation pairs with strong dominance show weak epistasis and vice versa.

### Folding and binding mutants differ in their epistasis but have the same dominance

When mutations are combined in the same allele in Model 2, the outcome differs depending on whether the mutations affect folding or binding (Figure 3c, e, f). That is, combining single mutants with the same phenotypic value but different underlying causal biophysical mechanisms results in a different outcome in a double mutant. This is because the non-linear relationships between the phenotype and the free energies of folding or binding are different (Supplementary Figure 3a, b) (Faure et al., 2022; Li and Lehner, 2020; Otwinowski, 2018).

In contrast, when two mutations in different alleles are combined, the phenotype of the double mutant does not depend on whether the mutations affect folding or binding (Figure 3d, g, h). Thus, in this simple folding and binding system, the magnitude of epistasis but not dominance depends on the biophysical effect of a mutation – whether a mutation affects protein stability or the binding affinity.

### Changes in ligand concentration switch alleles from dominant to recessive

In the version of Model 2 presented in Figure 3, the ligand concentration is the same as the total protein concentration. However, as illustrated in Figure 4, altering the ligand concentration both quantitatively and qualitatively changes the dominance.

**Figure. 4.**
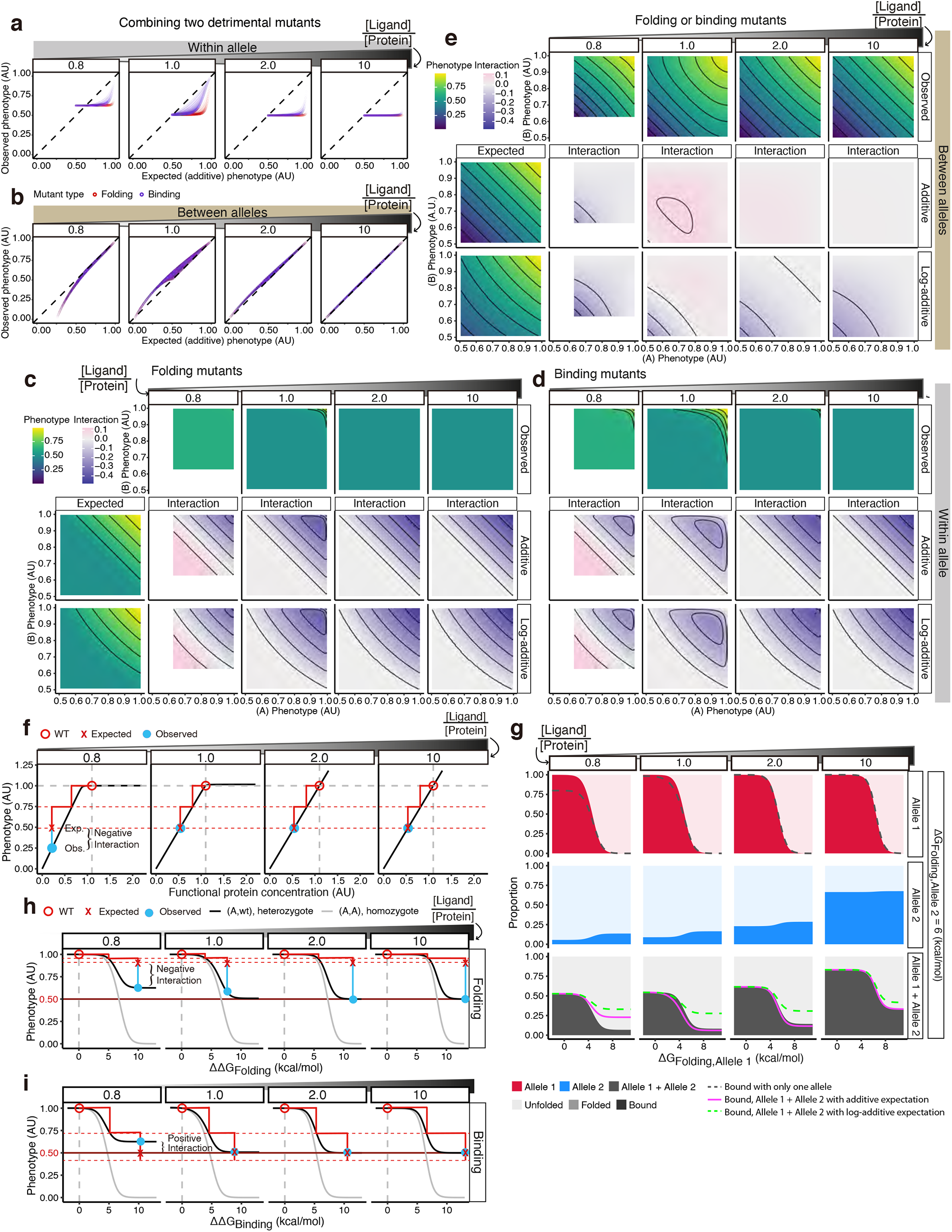
Changes in ligand concentration switch alleles from dominant to recessive. (a, b) The relationships between the observed and expected phenotypes with additive expectation when combining two detrimental mutants within- (a) or between-allele (b) at different ligand-protein ratios. (c-e) Heatmaps show how two mutations combine within (c, d) or between alleles (e) when they both affect the same biophysical parameters: protein-folding (c, e) or ligand-binding (d, e). (f) The relationship between the functional protein concentration and the relative proportion of the phenotype at different ligand-protein ratios. The circles and red lines show how two mutations of the same phenotype (0.75 AU) are expected or observed to combine based on the dose-response curves in the model. (g) Proportion of ligand-bound, folded but unbound, and unfolded proteins in each cell with Allele 2 protein folding energy = 6 kcal per mol while Allele 1 protein folding energy changes. (h, i) The relationship between the change in free energy of protein folding (h) or ligand-binding energy and phenotype at different ligand-protein ratios. The circles and red lines show how two mutations of the same phenotype (0.75 AU) are expected or observed to combine.

At high ligand concentrations (e.g. ligand:protein = 10 in Figure 4b), there is no dominance, with mutations combining additively. When the ligand concentration is reduced (e.g. ligand:protein = 2 in Figure 4b), the phenotype of a double mutant is better than the additive expectation and this tendency further increases at ligand:protein = 1 (Figure 4b). However, as the ligand concentration is further decreased, the phenotypes of double mutants become worse than the additive expectation, i.e., the more detrimental allele changes from recessive to dominant (ligand:protein = 0.8 in Figure 4b). Thus, changing the concentration of a ligand both quantitatively and qualitatively alters the interactions between alleles, resulting in the more detrimental variant switching from recessive to dominant.

Why does ligand-binding cause dominance and why does this dominance switch from positive to negative as the ligand concentration changes? When the concentration of the ligand is in excess, changes in protein concentration result in a proportional change in ligand binding and the two alleles effectively behave as independent thermodynamic systems (Figure 4f, g, Supplementary Figure 4). However, when the ligand concentration is reduced, the two alleles now compete for binding to the ligand and so they can no longer be considered as independent systems. When a mutation A destabilizes allele 1, less of allele 1 binds to the ligand but, because ligand binding is competitive, more of the ligand now binds to allele 2, resulting in a smaller than additive reduction in the total protein-ligand complex (Figure 4g, Supplementary Figure 4). However, as the ligand concentration is further reduced, the system enters a regime where the relationship between the fraction bound and the total protein concentration is no longer linear as the protein is in excess and all of the ligand is bound (Figure 4f, Supplementary Figure 4). Moderately reduced stability or affinity now has no effect on the concentration of the protein-ligand complex such that only larger changes in energy alter the concentration of the bound complex (Figure 4f, g, Supplementary Figure 4). As a result, many detrimental mutations combine to have greater than additive effects (Figure 4e).

### Changes in ligand concentration alter epistasis

The interactions between mutations within the same allele also change as the ligand concentration is altered with the appearance of positive epistasis at low ligand concentrations (Figure 4a, c, d). For example, when the ligand protein ratio is 0.8, 80% of the ligand is bound when both alleles are wild-type but 50% of the ligand is bound when one allele is null. The lowest possible phenotypic value is thus 62.5% of the wild-type phenotype (0.5/0.8) not 50% of the wild-type as the additive and log-additive models assume (Figure 4h, i).

To sum up, variations in the ligand concentration alter epistasis quantitatively when two mildly detrimental mutations combine (Figure 4h) and qualitatively when two very detrimental mutations combine (Figure 4i).

### Nonlinear dose-response curves differentially transform dominance and epistasis

So far, we have considered situations where a phenotype is linearly dependent on the concentration of a folded protein or a protein-ligand complex. However, in reality, phenotypes often depend nonlinearly on the concentrations of macromolecules (Keren et al., 2016). We therefore used three representative concentration-phenotype relationships – concave, convex and sigmoidal linking functions – to explore how non-linear relationships alter the interactions within and between alleles (Figure 5a).

**Figure. 5.**
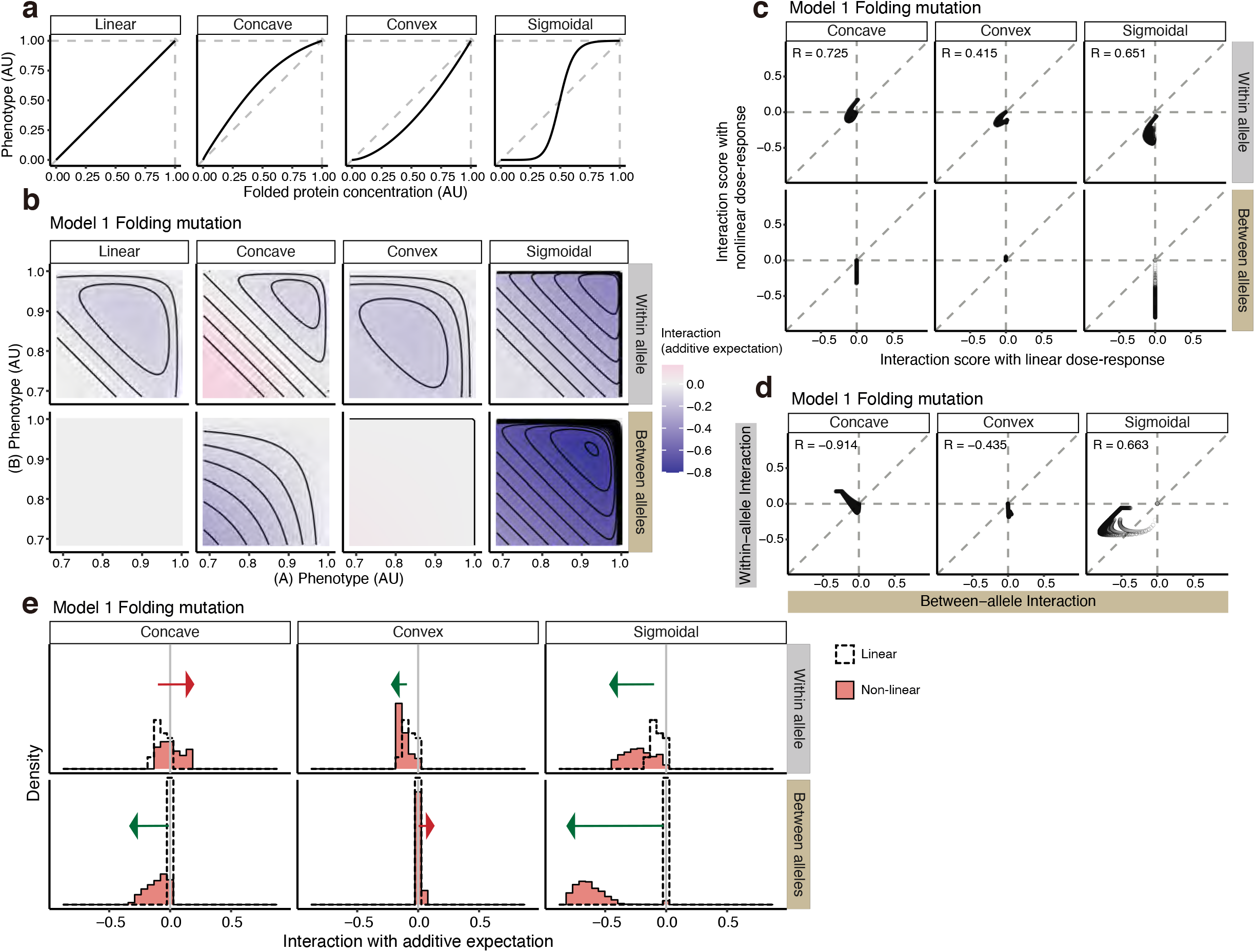
Nonlinear concentration-phenotype functions differentially transform dominance and epistasis. (a) Linear, concave, convex and sigmoidal linking functions used to transform protein concentrations to phenotypes. (b) Interaction scores based on the additive expectation for double mutants within- or between-allele with linear (Model 1), concave, convex, or sigmoidal protein concentration – phenotype relationships. (c) With vs. without nonlinear linking function comparisons of interaction scores based on the additive expectation. (d) Between- vs. within-allele double mutants interaction scores based on the additive expectation, with nonlinear linking functions. (e) Distribution of interaction scores based on additive expectation before and after nonlinear linking functions. The green arrow indicates the distribution shifting towards negative values while the red arrow indicates the distribution shifting towards positive values; the arrowheads point at the range after applying the nonlinear linking functions to the phenotype.

When applied to the protein-folding model (Model 1), all three functions alter both within- and between-allele genetic interactions (Figure 5b, c). Moreover, the interactions not only change quantitatively but also in some cases qualitatively, switching from positive to negative (Figure 5c). The effects of the non-linear concentration fitness functions can also differ for within- and between-allele interactions. For instance, a concave function shifts within-allele interactions to less negative or positive values, whereas between-allele interactions become negative (Figure 5c, e). In contrast, a sigmoidal concentration-fitness function shifts both within- and between-allele interactions towards more negative values, with a stronger effect on the latter (Figure 5c, e). Moreover, between- and within-allele genetic interactions can become anti-correlated when nonlinear linking functions are applied (Figure 5d, compare to Figure 3i, j). The effects of linking functions on the interactions between variants in Model 2 are shown in Supplementary Figure 5.

Taken together, these results show that protein folding generates epistasis and a single binding reaction is sufficient to generate dominance, meaning that epistasis and dominance should be widely expected. However, the strength and sign of epistasis and dominance are highly dependent on the details of the system, for example, on the relationship between the concentration of a molecule and the phenotype of interest, making them difficult to predict without a detailed mechanistic understanding of a system and knowledge of the relevant cellular parameters or large-scale empirical measurements of mutational effects and interactions. For example, even the simple perturbation of altering the concentration of a binding partner can quantitatively alter epistasis and switch an allele from dominant to recessive.

## Discussion

We have analyzed here how mutations interact in simple thermodynamic models of protein folding and binding to better understand the origins and expectations for the interactions between mutations within and between the alleles of a gene: epistasis and dominance, respectively. Protein folding alone generates epistasis and the addition of a single binding reaction is sufficient to generate dominance. How two mutations interact often differs in both magnitude and sign depending upon whether they combine within the same allele or in different alleles. Moreover, interactions depend qualitatively and quantitatively on both the biophysical effects of mutations and the context, with, for example, a change in the concentration of a ligand sufficient to switch a mutation from dominant to recessive in even the simple system of a protein that folds and binds a single ligand.

Taken together, our results illustrate that both dominance and epistasis should be widely expected in even the simplest biological systems, but also emphasize that they are system properties and therefore difficult to predict without detailed mechanistic understanding and parameter measurements or, alternatively, extensive empirical quantification of mutational effects and interactions in a system.

That dominance and epistasis are (1) expected in even the simplest systems, (2) context-dependent, and (3) difficult to predict – has important implications for human genetics, biotechnology and evolution. As an example, our results suggest that it is not unreasonable to expect disease-causing alleles to switch from dominant to recessive depending upon the conditions, examined phenotype or individual.

An important simplification of our approach is that it assumes that free energy changes are additive. Although this is likely to be true for the majority of mutation combinations (Faure et al., 2022; Otwinowski, 2018), specific non-additive changes in free energy, for example between mutations in physically contacting residues (Carter et al., 1984; Horovitz et al., 2019), will generate additional epistasis and dominance.

In future work, it will be interesting to quantify and compare the expected dominance and epistasis in more complex equilibrium systems, as well as in dynamical models (Gjuvsland et al., 2013; Omholt et al., 2000). In addition, it will be important to experimentally evaluate dominance and epistasis in the same genes, for example, using deep mutational scanning approaches (Domingo et al., 2019). Using large-scale mutagenesis approaches it will also be possible to evaluate how frequently specific exceptions to the typical patterns of dominance and epistasis arise, what the most frequent causes of these exceptions are, and how they can be predicted.

An important general goal in human genetics, agricultural genetics, biotechnology and evolutionary biology is to improve genetic prediction beyond the accuracy that can be achieved with additive models. Here we have shown that, even in the very simplest biological systems, epistasis and dominance are expected, context-dependent and challenging to predict. It is precisely the emergent, plastic and difficult-to-predict nature of dominance and epistasis that makes better-than-additive genetic prediction a formidable challenge.

How in practice will better-than-additive genetic prediction be achieved? One approach could be to build detailed mechanistic models of each system of interest and to parameterize these models using experimental measurements. However, given the complexity of most biological systems of interest, we suggest that a non-mechanistic approach may actually prove more successful. Indeed, we suspect that the combination of large-scale data collection and machine learning may prove to be the more efficient strategy for improving genetic prediction beyond what can be achieved with additive models and that this will accelerate progress in biotechnology, agriculture and medicine.

## Materials and methods

### Phenotypes

To study how mutations in a protein-coding gene combine in a diploid system, we first defined the phenotype (W) in the system as functional molecule concentration inside the cell relative to the wild-type diploid situation. These functional molecules can be folded proteins or a protein-ligand complex.

In each model, the phenotype of a mutant is normalized to the homozygous wild-type (wt) phenotypes so that W_wt,wt_ = 1 AU and the complete loss of function is set to 0. With the simple assumption of additivity in the functional molecule concentrations, we imposed a lower bound of the expected simple heterozygote phenotype (i.e. any single mutants or any double mutants’ expected phenotypes within the same allele) to 0.5 AU (0.5X of the wild-type phenotype) (Figure 1d).

### Expected phenotypes and interaction scores

When there is only one mutation A, the homozygote is annotated as (A,A) and the heterozygote with the wild-type is annotated as (A,wt). The compound heterozygote is annotated as (A,B) while the heterozygous double mutant with both variants present in the same allele is designated (AB,wt).

For a given pair of single mutant phenotypes, there are two expected double mutant phenotypes based on additive and log-additive expectations as shown in the following equations Eq. (1, 2).

Additive expectation:

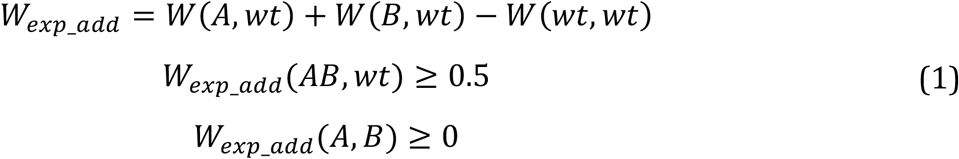

Log-additive expectation:

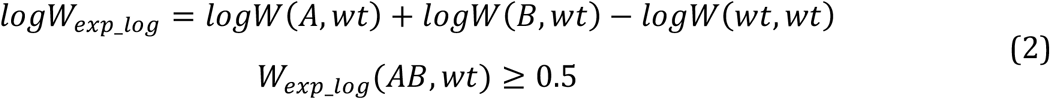

For within- and between-allele interactions, the only differences in the expected phenotypes are the lower limit, that is, within-allele mutation combinations are set to 0.5 while the between-allele limits are 0 as shown in Eq. (1, 2).

Interactions between mutations are quantified as differences between observed versus expected phenotypes for both within- and between-allele interactions and regardless of the additive model used (additive or log-additive), as follows:

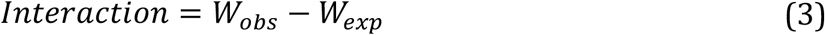

### Model 1: Protein folding

Phenotype is determined by the total concentration of folded protein. In this model, the protein of interest (X) expressed from each allele (allele α i, I ∈{1, 2}) has two configuration states: unfolded (X_U, α i_) and folded (X_F, α i_). The free energy difference between folded and unfolded protein states is ΔG_Folding, α i_ (kcal per mol). Mutations on each allele can affect folding energy (ΔG_Folding_), which is described as the sum of wild-type folding energy and the energy differences (mutations) ΔG_Folding,wt_ + ΔΔG_Folding, α i_. Equilibrium between the two states follows Eq. (4).

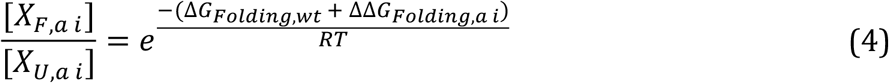

In the above and following equations, R is the gas constant (R = 1.98 × 10^−3^ kcal per mol), T is the absolute temperature for 37 °C (310.15 Kelvin) and the wild-type ΔG_Folding,wt_ is set to −2 kcal per mol unless stated otherwise.

The total concentration of the protein (X_T_) follows Eq. (5).

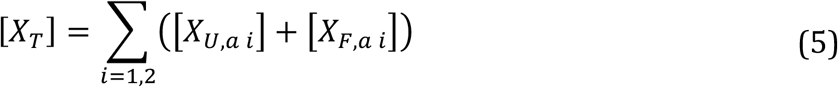

Expression levels from each allele were considered to be equal and therefore, [X_T,1_] = [X_T,2_] = 0.5 [X_T_] in all our models. Using Eq. (4) and (5) with [X_T_] as a constant, we can calculate the functional molecule [X_F,α i_] as a function of energy terms and total protein concentration in the following way:

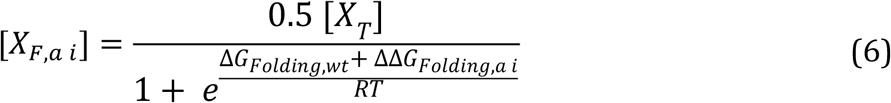

The phenotype of a mutant W_mut_ is the sum of the folded protein concentration normalized to that of the wild-type, following Eq. (7):

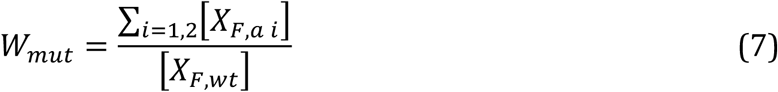

To obtain the observed double mutant phenotypes between alleles, we could combine Eq. (4 - 7) and simply replace ΔΔG_Folding, α 1_ with ΔΔG_Folding,A_ and ΔΔG_Folding, α 2_ with ΔΔG_Folding,B_ respectively, as shown below.

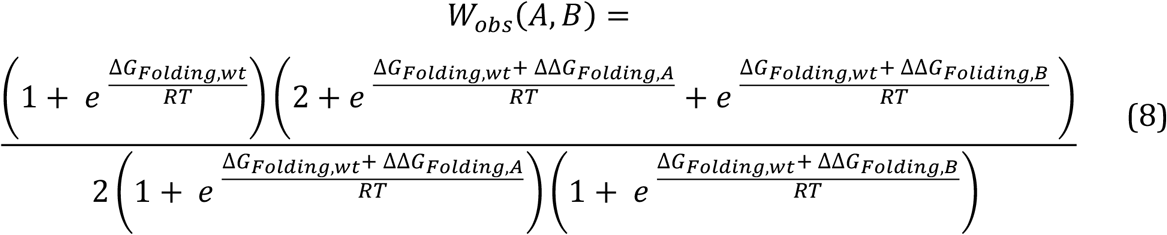

For a single mutation A, we simply set ΔΔG_Folding, α 2_ = 0 and replace ΔΔG_Folding, α 1_ with ΔΔG_Folding,A_ in Eq. (8). After simplification, the equation for a single mutant phenotype can be written as follows:

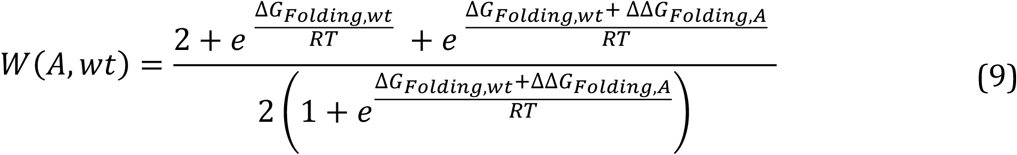

For the double mutation (A and B) in the same molecule, ΔΔG_Folding,A_ and ΔΔG_Folding,B_ values are added to replace ΔΔG_Folding,A_ in Eq. (9), resulting in Eq. (10).

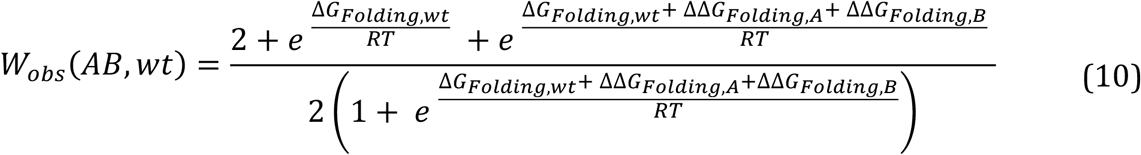

Additive or log-additive expectations of the double mutant phenotypes are calculated by combining Eq. (1) or Eq. (2) with Eq. (9) for both within-allele and between-allele combinations.

We note that the additive expectation for between-allele mutation combinations is mathematically the same as observed between-allele mutation combinations (Figure 2i, k, upper right). Firstly, by combining Eq. (1) with Eq. (9) for each mutation, we obtain Eq. (11):

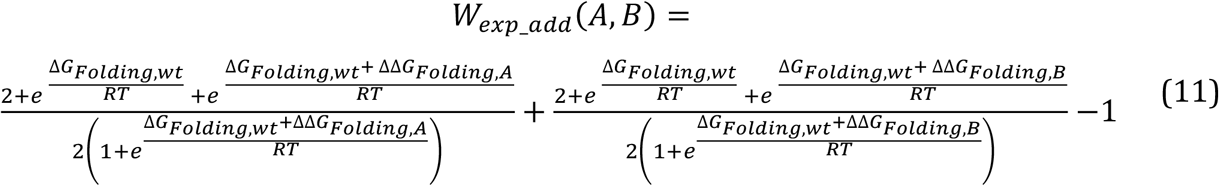

Then, Eq. (11) can be rewritten into the same form as Eq. (8). In other words, mathematically W_exp_add_(A,B) = W_obs_(A,B) in this model. The log-additive expectation is, however, different.

### Model 2: Folding and Ligand-binding

Phenotype is determined by the total concentration of protein-ligand complex in Model 2. In this model, one protein molecule binds to one ligand molecule and the total ligand concentration (L_T_) is the sum of free ligand (L) and those bound to the protein (equal to the protein-ligand complex concentration, ∑_i=1,2_(X_L, α i_)), which follows Eq. (12).

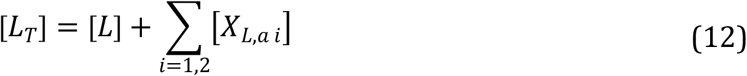

Compared to Model 1, there is an additional energy term – the free energy difference between the protein-ligand complex state and the unbound folded protein with free ligand state ΔG_Binding_ (kcal per mol) – which we define as binding energy. Mutations on each allele can now be described as those affecting protein folding energy (ΔΔG_Folding_) or binding energy (ΔΔG_Binding_).

There are three configuration states of the protein expressed from each allele: unfolded (X_U, α i_), folded (X_F, α i_) and protein-ligand complex (X_L, α i_). The equilibrium between [X_U, α i_] and [X_F, α i_] follows Eq. (4), and the equilibrium between [X_F, α i_] and [X_L, α i_] follows Eq. (13).

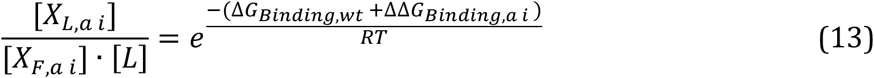

The total concentration of the protein (X_T_) follows Eq. (14), as shown below.

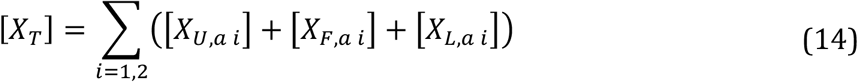

As stated earlier, [X_T, α 1_] = [X_T, α 2_] = 0.5 [X_T_]. By combining this information with Eq. (4), Eq. (12-14), we express the protein-ligand complex concentration from each allele as a function of energy terms, total protein concentration and the free ligand concentration as shown in Eq. (15):

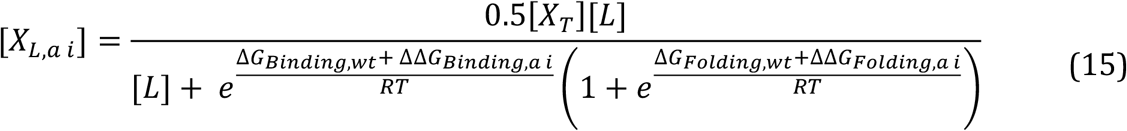

Phenotype of a mutant (W_Mut_) is the sum of the protein-ligand complex concentration normalized to that of the wild-type as shown below in Eq. (16).

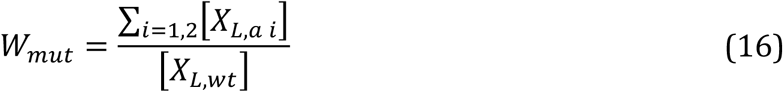

For a double mutant (A and B mutations respectively) between alleles, we combine Eq. (4, 12-16) and simply replace α 1 with A and α 2 with B for corresponding mutation types, as shown in Eq. (17).

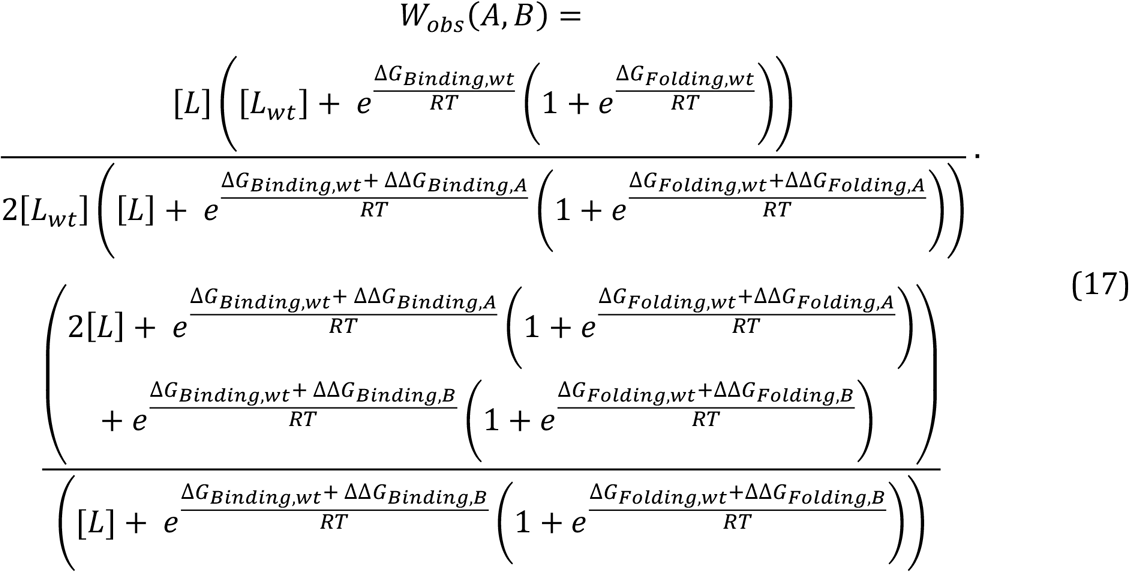

If there is only one mutation A on allele 1, then ΔΔG_Folding, α 2_ = ΔΔG_Binding, α 2_ = 0 and we replace ΔΔG_Folding, α 1_ with ΔΔG_Folding,A_ and ΔΔG_Binding, α 1_ with ΔΔG_Binding,A_. Eq. (17) will be simplified to a single mutant phenotype as follows:

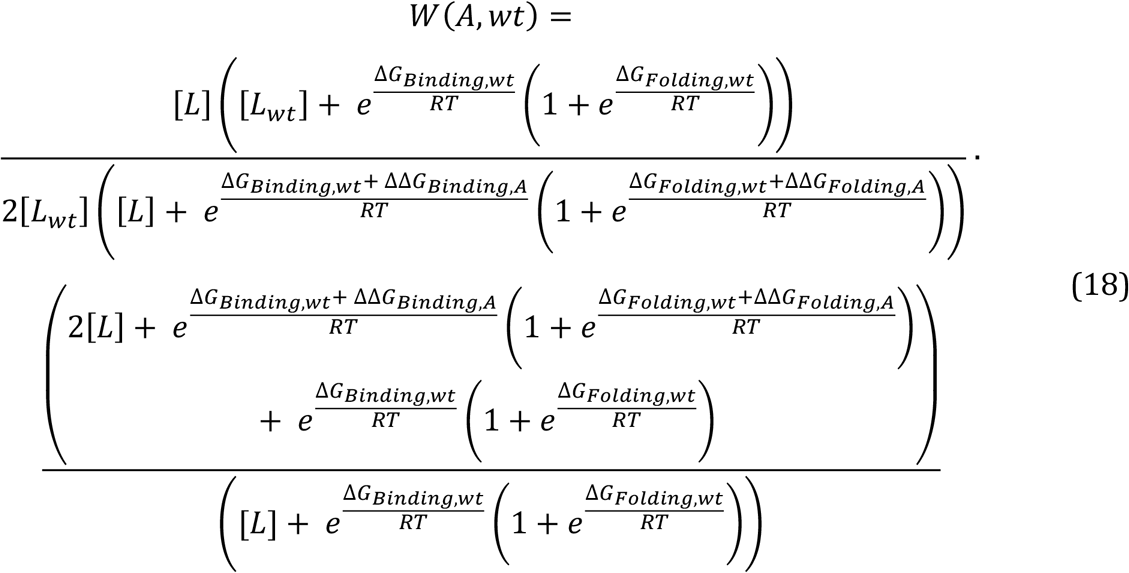

Above, [L_wt_] is the free ligand concentration inside the wild-type while [L] is the free ligand concentration inside the mutant cells. For a double mutant on the same allele, ΔΔG_Folding,A_ and ΔΔG_Folding,B_ are summed in place of ΔΔG_Folding,A_, and ΔΔG_Binding,A_; ΔΔG_Binding,B_ are summed in place of ΔΔG_Binding,A_ while keeping allele 2 as wild-type (ΔΔG_Folding, α 2_ = 0 and ΔΔG_Binding, α 2_ = 0), shown as in Eq. (19).

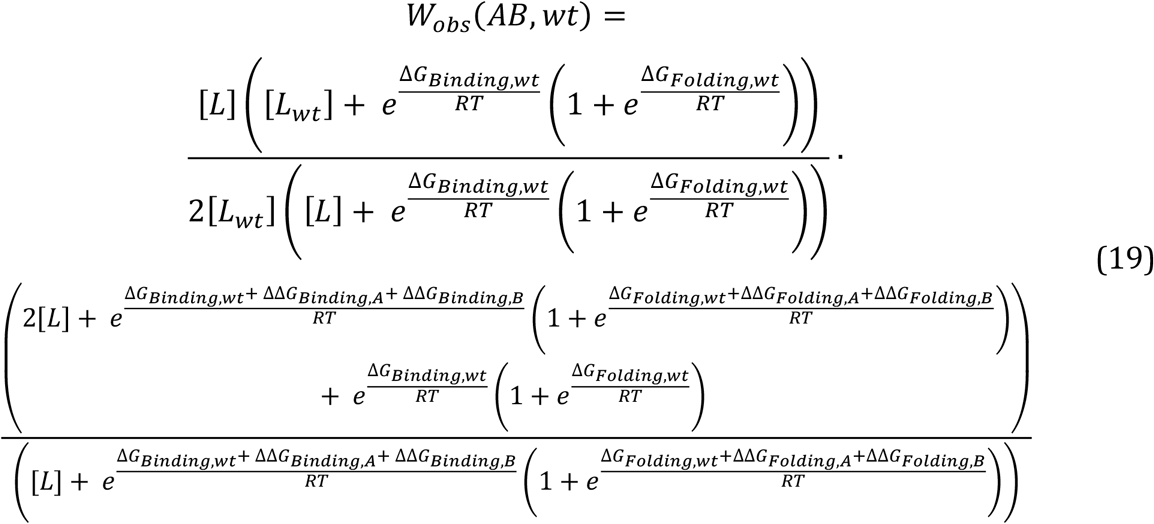

Additive or log-additive expectation can be calculated by combining Eq. (1) or Eq. (2) with Eq. (18), with corresponding parameters changed in the equations.

To be noted, the free ligand concentrations in the cell will change with the change of corresponding phenotypes. For example, the free ligand concentrations in single mutant A or B are [L_A_] = [L_T_] - ([X_L,A_] + [X_L,WT_]) or [L_B_] = [L_T_] - ([X_L,B_] + [X_L,WT_]). If [L_T_] is much bigger than [X_T_] and thus the free ligand concentration is not affected by how many molecules are bound to the protein, free ligand concentration can be considered a constant as [L_T_] (i.e. [L_A_] ≈ [L_B_] ≈ [L_T_]). In this situation, the additive expectation for a between-allele mutant combination simplifies to Eq. (17), indicating that there will be no dominance when the total ligand concentration is much bigger than the total protein concentration.

We examined several cases where we altered [L_T_] so that the total ligand concentration is the same, more abundant (2X, 10X) or less abundant (0.8X) than the total protein concentration [X_T_]. With mutational effects as changes in folding energy (ΔΔG_Folding, α i_) or ligand-binding energy (ΔΔG_Binding, α i_) as input, we calculated the phenotypes using the R package rootSolve.

### General non-linear curves

Three common types of non-linear functions (concave, convex, and sigmoidal) linking functional protein concentrations to the downstream phenotypes were used. All the curves go through the points (0,0) and (1,1). In the equations below, x and y define phenotypes before and after applying each of the non-linear transformations.

Concave curve:

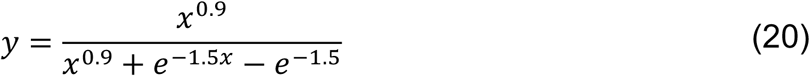

Convex curve:

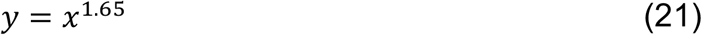

Sigmoidal curve:

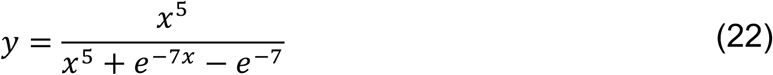

### Simulation of mutational effects and solving equations

Wild-type biophysical parameters and relative ligand concentration to the total protein concentrations in each model are shown in the table below (Table 1).

**Table 1.** Parameter values

| | Mutation types | $\Delta G_{\text{Folding,wt}}$<br>(kcal per mol) | $\Delta G_{\text{Binding,wt}}$<br>(kcal per mol) | $L_T$<br>Ratio to $X_T$ |
| --- | --- | --- | --- | --- |
| Model 1 | Folding | -2/-3.5/-0.5 | NA | NA |
| Model 2 | Folding,<br>Binding | -2 | -5 | 1/0.8/2/10 |

F: Folding. L: Ligand-binding. “/” separates different parameter options that we examined in this study and NA indicates that the parameter is not relevant in the model.

Two mutation types – those affecting protein stability (folding mutations) and binding affinity to a ligand (binding mutations) were described as changes in Gibbs free energy (ΔΔG) between folded-unfolded states (ΔΔG_Folding_) and bound-unbound states (ΔΔG_Binding_). To generate single mutations, ΔΔG_Folding_ and ΔΔG_Binding_ ranging from −2 to 13 kcal per mol with an interval of 0.125 kcal per mol were added to ΔG_Folding,wt_ and ΔG_Binding,wt_ respectively.

To examine how two mutations of given phenotypes combine, we generated single mutants with phenotypes ranging from 0.5 AU to 1.02 AU, with an interval of 0.005 AU. Using the same sets of equations, and phenotypes as inputs, again using R and rootSolve package, we calculated the corresponding ΔΔG for each phenotype. Then, with the obtained ΔΔG values for each mutant as new inputs to the system, we calculated double mutants’ phenotypes.

As stated earlier, when two mutations affecting the same biophysical parameters combined on the same allele, ΔΔG was added to the corresponding parameter as the new input to calculate double mutants’ phenotypes. On the other hand, when mutations combined between alleles, ΔΔG was kept separate and inputted as two independent values on each allele for the phenotype calculation.

The processes of calculating phenotypes from mutants (ΔΔG) and ΔΔG from phenotypes with the nonlinear linking functions are the same as those without linking curves, except that the phenotypes were transformed based on the Eq. (20 - 22) depending on the situation. The transformed single mutant phenotypes were used to calculate additive or log-additive expectations based on Eq. (1, 2), and interaction scores were calculated with Eq. (3). All the phenotype and interaction patterns were visualized using ggplot package in R.

## Code availability

https://github.com/XLi-Lab/P1_Dominance_vs_Epistasis

## Acknowledgments

X.X. performed all analyses and made the figures with the assistance of X.L.. X.L. and B.L. conceived the project, designed the analyses and wrote the manuscript with input from X.X..

Work in the lab of X.L. is supported by the Young Scientists Fund of the National Natural Science Foundation of China (NSFC Grant No. 32100478). Work in the lab of B.L. is supported by the European Research Council (ERC, Advanced Grant ‘Mutanomics’ 883742), the la Caixa Research Foundation (LCF/PR/HR21/52410004), the Bettencourt Schueller Foundation, the AXA Research Foundation, the Spanish Ministry of Science, Innovation and Universities (PID2020-118723GB-I00), Agencia de Gestio d’Ajuts Universitaris i de Recerca (AGAUR, 2017 SGR 1322) and the CERCA Program/Generalitat de Catalunya. We also acknowledge the support of the Spanish Ministry of Science and Innovation to the EMBL partnership and the Centro de Excelencia Severo Ochoa.

---

**Supplementary Figure 1.**
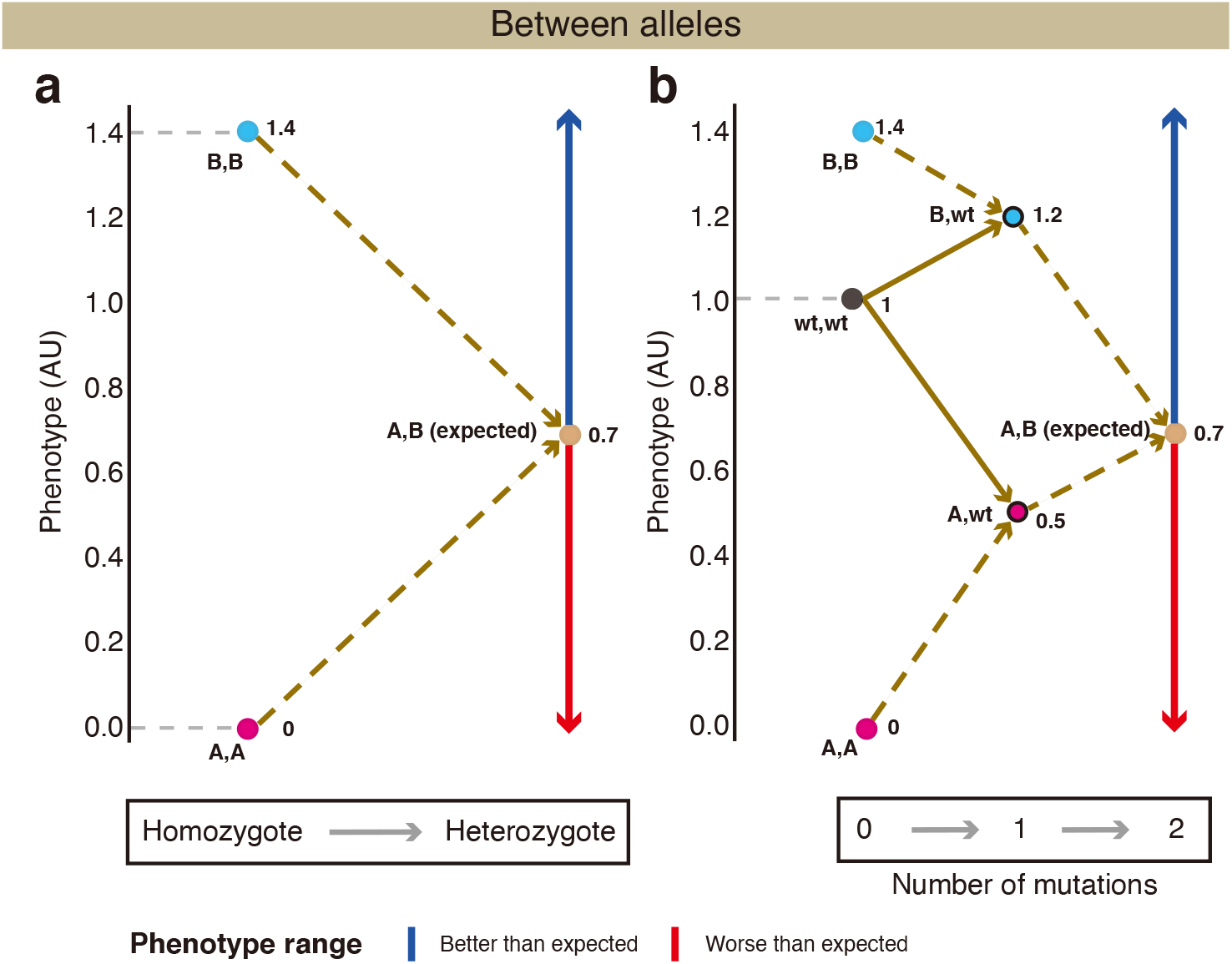
Comparison of two homozygous combinations and two corresponding heterozygous combinations. (a, b) The scheme of quantifying how two homozygotes combine (a) and how two corresponding heterozygotes combine (b) between alleles with the additive assumption.

**Supplementary Figure 2.**
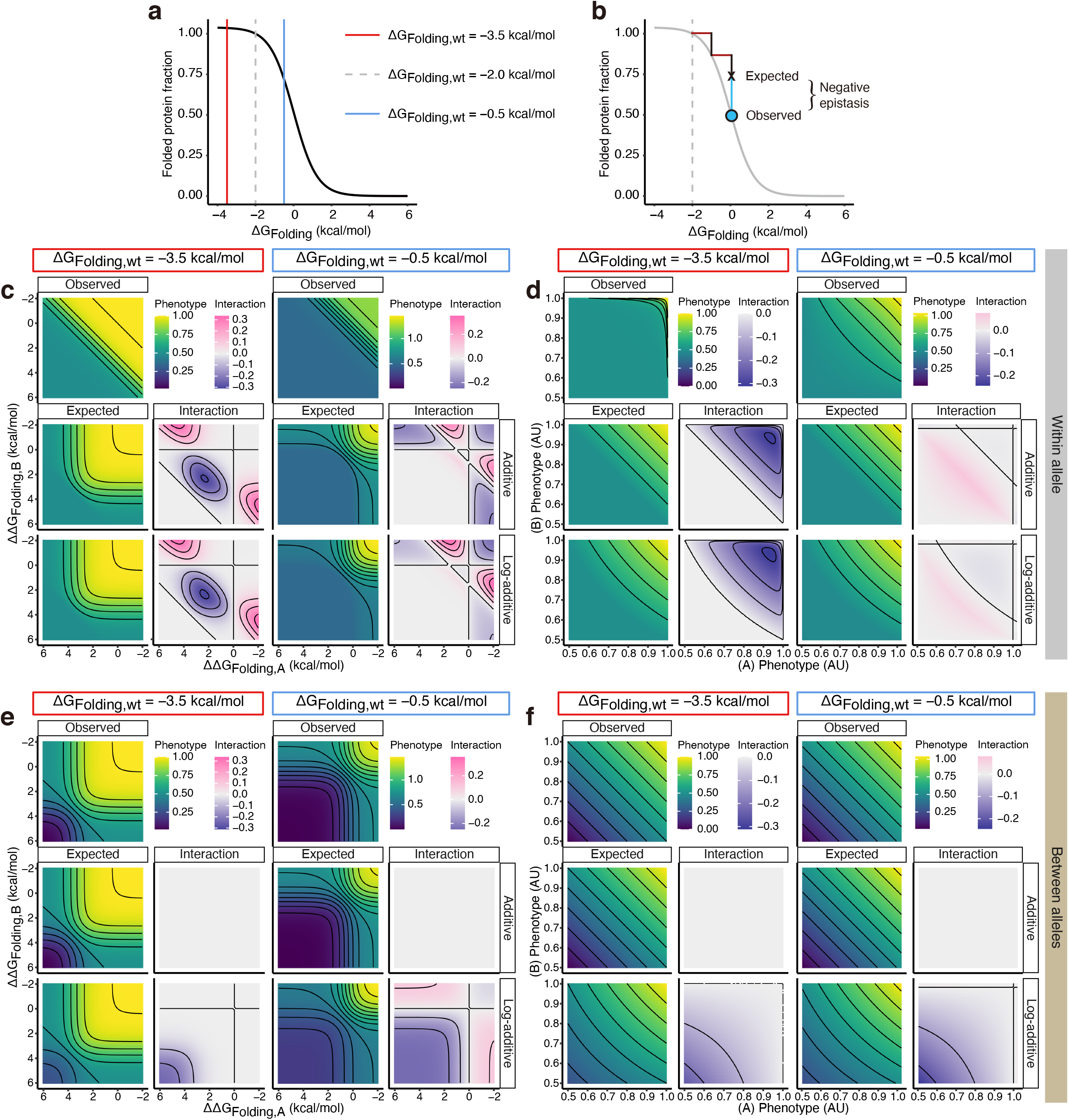
Protein folding generates epistasis but not dominance. (a) Relationship between the free energy of protein folding and folded protein fraction. Vertical lines indicate three different settings of the wild-type stability with the dashed line showing the default setting presented in Figure 2. (b) Non-specific epistasis arises from the nonlinear relationship between free energy and phenotype. (c-f) Heatmaps of how two mutations combine within- (c, d) or between-allele (e, f) ordered by free energy changes (c, e) or phenotypes (d, f) for a very stable protein (the red box title) or marginally stable protein (the light blue box title).

**Supplementary Figure 3.**
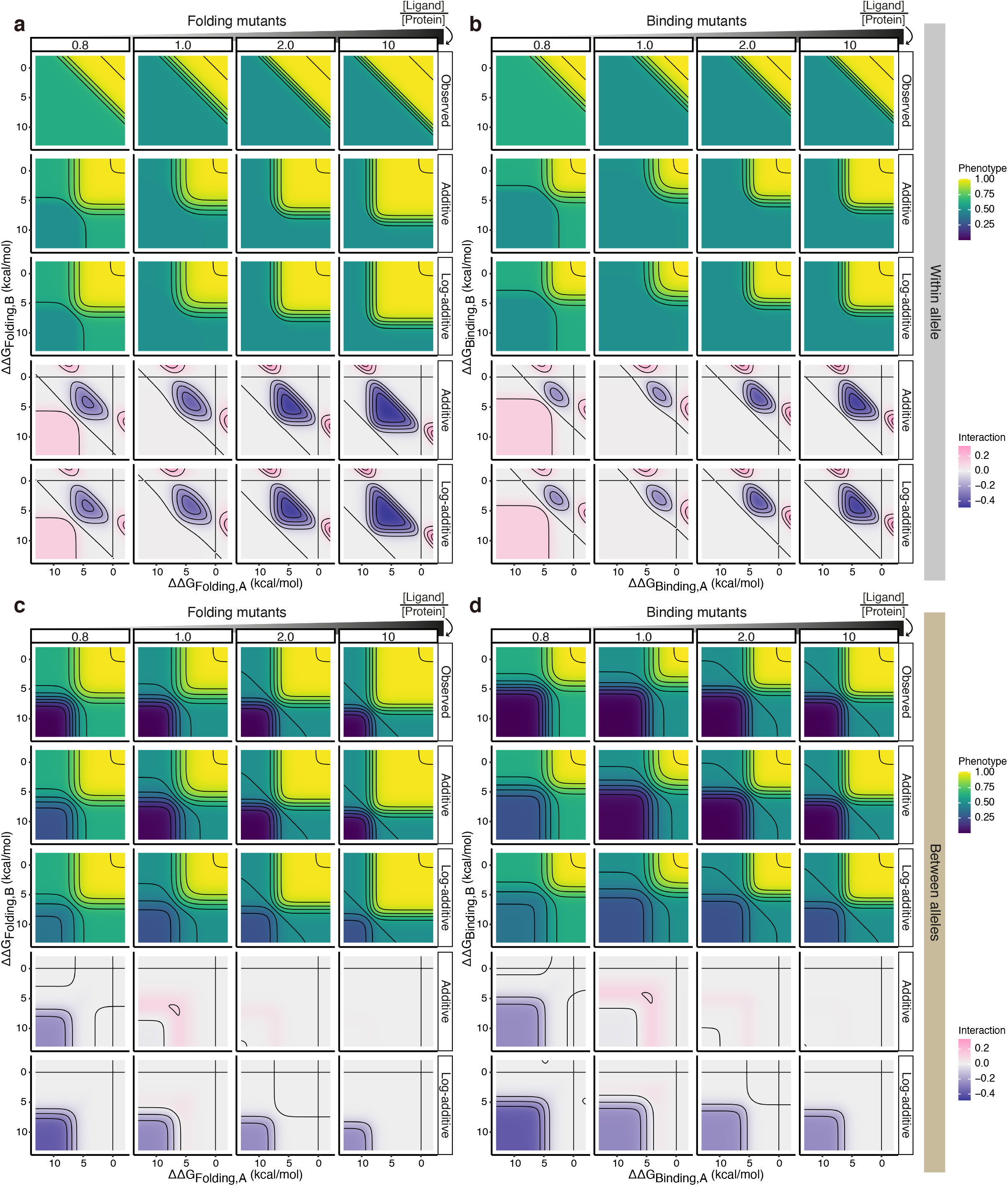
Ligand binding generates dominance. (a-d) Heatmaps show how two mutations combine within (a, b) or between alleles (c, d) when they both affect the same biophysical parameters: protein-folding (a, c) or ligand-binding (b, d) at different ligand-protein ratios.

**Supplementary Figure 4.**
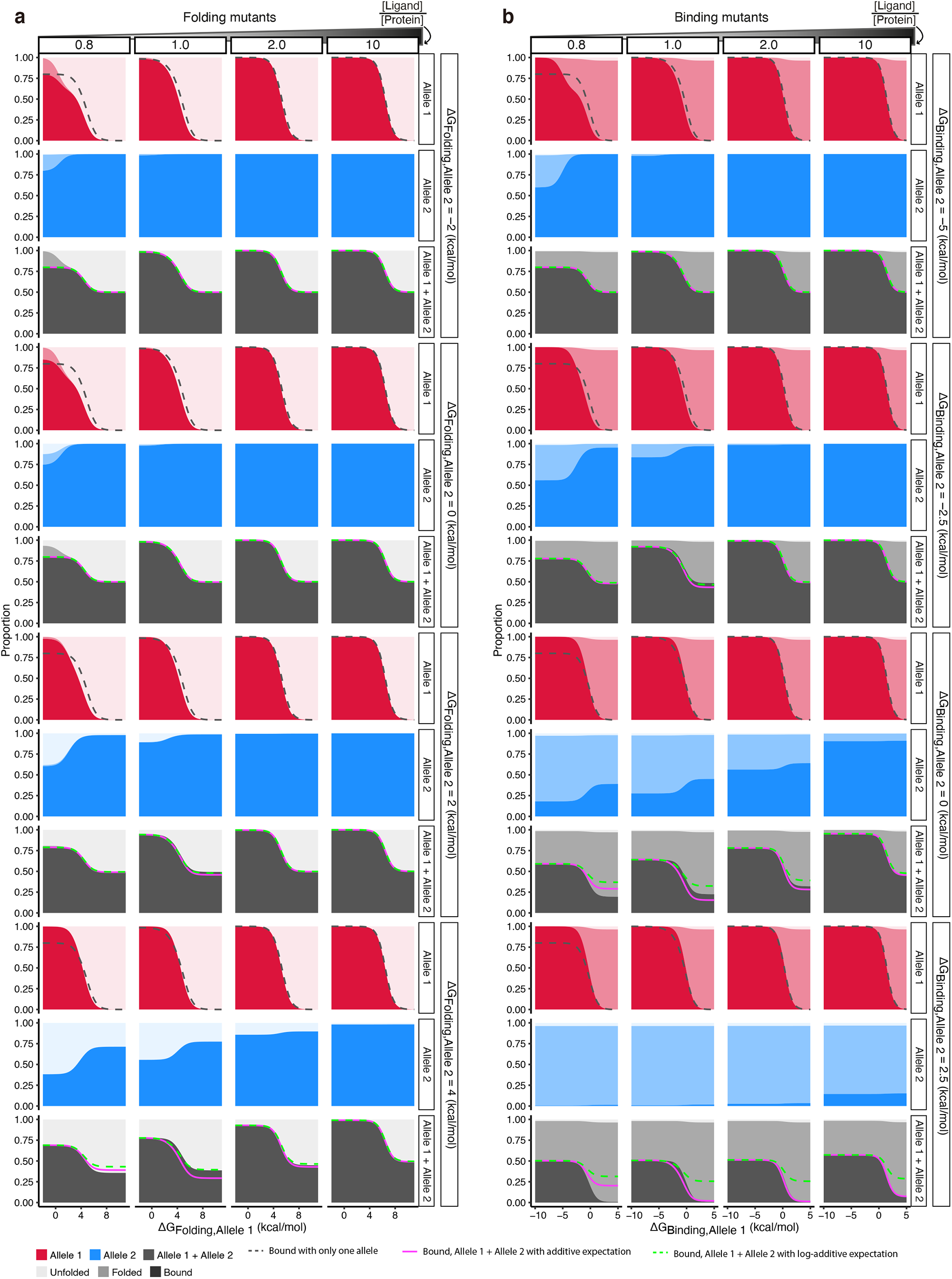
The proportion of protein states while altering Allele 1 energy at differing ligand concentrations. Allele 1 and Allele 2 protein partitioning when the mutations alter Allele 1 protein folding energy (a) or binding energy to the ligand (b), based on Model 2.

**Supplementary Figure 5.**
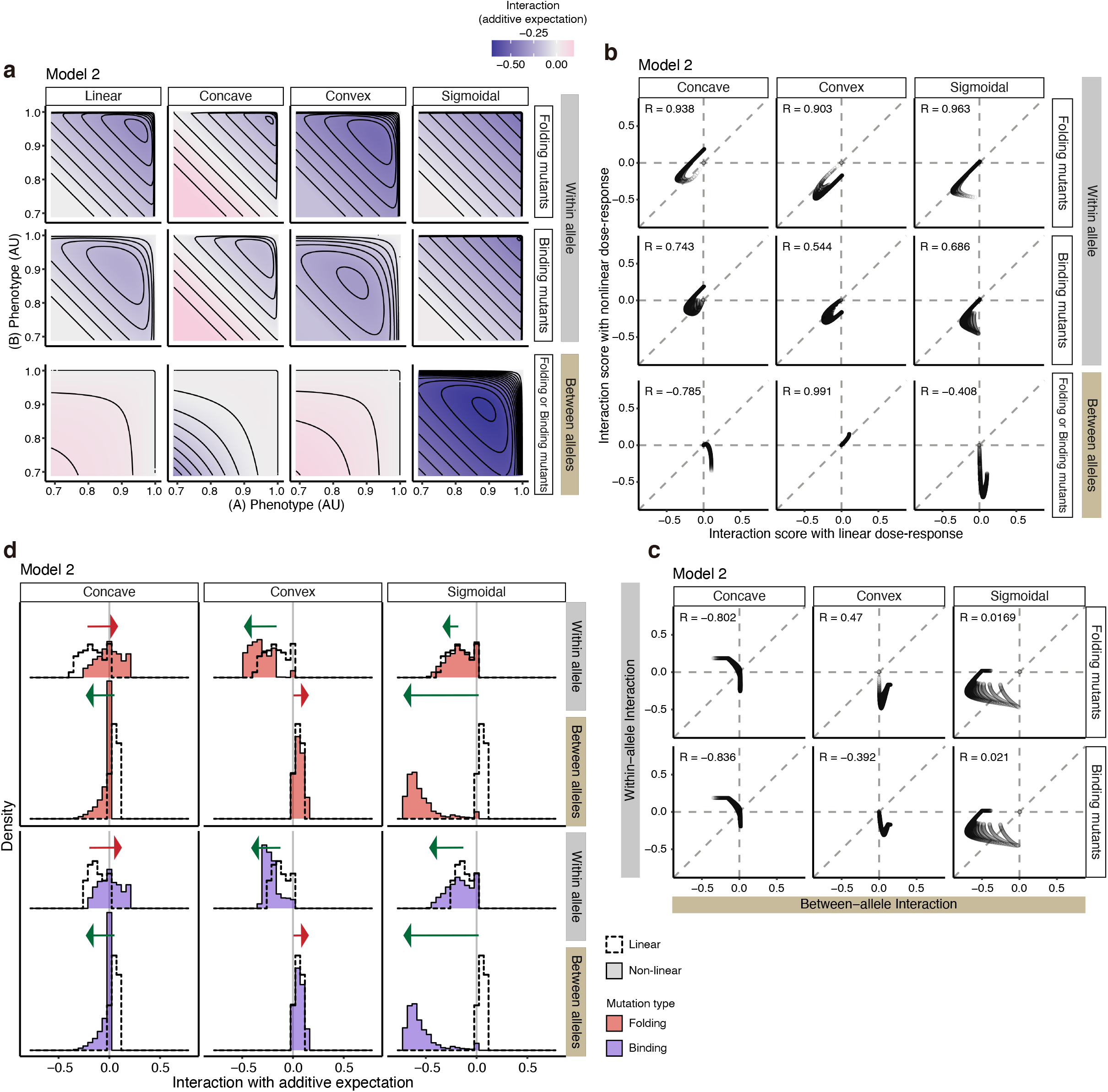
Nonlinear concentration-phenotype functions differentially transform dominance and epistasis in Model 2. (a) Interaction scores of double mutants within- (the two top rows) or between-allele (the bottom row) with linear, concave, convex, or sigmoidal protein concentration – phenotype relationships. (b) Comparisons of interaction scores between double mutants within- (the two top rows) or between-allele (the bottom row) with or without nonlinear linking functions. (c) Comparisons of between- vs within-allele interaction scores of double folding (the top row) or binding (the bottom row) mutants with nonlinear linking functions. (d) Interaction score distributions before and after nonlinear linking functions. The green arrow indicates the distribution shifting towards negative values while the red arrow indicates the distribution shifting towards positive values. The arrowheads point at the range after applying the nonlinear linking functions to the phenotype.

